# The DC1 domain protein Vacuoleless Gametophytes positively regulates salt stress tolerance in *Arabidopsis thaliana*

**DOI:** 10.64898/2026.05.26.727883

**Authors:** Natalia Loreley Amigo, Maria Fernanda Marchetti, Salvador Colombo Lorenzani, Leonardo Agustin Arias, Juan Ignacio Poo, Milagros Escoriza, María Elisa Picco, María Cecilia Terrile, Diego Fernando Fiol

## Abstract

Vacuoleless Gametophytes (VLG) is a DC1 domain-containing protein initially characterized as essential for the development of both female and male gametophytes in *Arabidopsis thaliana*. In addition, VLG regulates stamen development through the involvement in lignin and jasmonic acid biosynthesis pathways. In this work, we report that VLG is also involved in salt stress tolerance in *A. thaliana*. Under salt stress, *VLG-*knock-down plants exhibited reduced germination, root elongation, biomass accumulation, photosynthetic pigment content, along with diminished expression of key salt-responsive genes. Conversely, these plants accumulated higher anthocyanins, and reactive oxygen species (H_2_O_2_ and O_2_^−^) compared to wilt type, indicating impaired oxidative stress control. In contrast, *VLG*-overexpressing plants showed a salt stress resistant phenotype with enhanced biomass and increased expression of salt-responsive genes under saline conditions. Together, these findings uncover an unexpected role for VLG as a positive regulator of salt tolerance, expanding the functional scope of DC1 domain proteins beyond reproductive development and providing new insights into plant mechanisms of abiotic stress resilience.

## 1 Introduction

Salt stress is a major environmental constraint affecting plant growth and agricultural productivity worldwide. High salinity disrupts cellular homeostasis by causing osmotic stress, ion toxicity, nutrient imbalance, and oxidative damage, thereby impairing plant growth and development (Munns and Tester 2008, Yang and Guo 2018, Van Zelm et al. 2020). To survive under saline conditions, plants activate a wide range of adaptive responses involving ion transport systems, redox balance, and transcriptional reprogramming of stress-responsive genes that collectively maintain cellular and metabolic homeostasis (Zhang and Shi 2013, Ismail and Horie 2017, Chen et al. 2022).

Divergent C1 (DC1) domain proteins are a family of plant-specific proteins implicated in different developmental and stress-related processes (Bhaskar et al. 2015, D’Ippólito et al. 2017, Arias et al. 2022, Amigo et al. 2025). Although their molecular functions are still poorly characterized, increasing evidence suggests that several DC1 proteins participate in plant responses to abiotic stress. In wheat, the DC1 domain-containing protein TaCHP was identified as a salt-responsive protein, and its overexpression increased salinity tolerance in *Arabidopsis thaliana* (Li et al. 2010). Likewise, the cotton DC1 protein GhCHR reduced Na^+^ accumulation and promoted root growth under saline conditions (Gao et al. 2016). More recently, members of the tomato SlCHP family and the sweet potato protein IbCHR10 were also associated with improved tolerance to salt and saline-alkaline stress (Li et al. 2023, Kang et al. 2024, Du et al. 2026). Despite these observations, the contribution of most DC1 proteins to stress adaptation remains largely unknown, particularly regarding how development-associated DC1 proteins contribute to stress adaptation.

Vacuoleless Gametophytes (VLG) is a DC1 domain-containing protein previously reported as essential for gametophyte development in *A. thaliana* (D’Ippólito et al. 2017). Loss of VLG function disrupts central vacuole formation during embryo sac and pollen development, causing early arrest of both gametophytes. VLG localizes to the endomembrane system and interacts with several proteins, including SNARE-associated proteins, lipases, and transcription factors, suggesting that it may participate in the assembly or regulation of protein complexes associated with membrane-related processes (D’Ippólito et al. 2017). In addition, VLG shows broad expression in sporophytic tissues, particularly in vascular tissues and expanding cells, pointing to functions beyond gametophyte development. Interestingly, some VLG-interacting proteins have been previously linked to abiotic stress responses. Among them, the lipase LTL1 enhances salt tolerance in yeast and plants (Naranjo et al. 2006), whereas RAP2.6L, an AP2/ERF transcription factor identified in our yeast two-hybrid screening, has been associated with responses to salinity and drought stress (Guo et al. 2005, Krishnaswamy et al. 2011). These observations raised the possibility that VLG could participate in coordinating adaptive responses to environmental stress.

In this work, we investigated the role of VLG in plant responses to salt stress. Our results show that VLG positively regulates salinity tolerance in *A. thaliana. VLG-*knock-down plants exhibited hypersensitivity to salt stress, reduced growth, altered ROS accumulation, and lower expression of salt-responsive genes. In contrast, *VLG*-overexpressing plants showed improved growth performance, enhanced ionic homeostasis, and increased expression of genes associated with salinity adaptation. Altogether, our findings identify VLG as a new component involved in plant salt stress responses and broaden the functional relevance of DC1 domain proteins beyond reproductive development.

## 2 Materials and Methods

### 2.1 Plant material and growth conditions

*VLG*-overexpressing (OE1 and OE2) and *VLG*-knock-down (L1, L2, and L3) lines were generated in *A. thaliana* ecotype Columbia (Col-0) (D’Ippólito et al. 2017, Amigo et al. 2025). *ProVLG:GUS* plants were reported in D’ippolito et al. (2017).

Plants were grown on soil (soil:vermiculite:perlite, 3:1:1) in growth chambers under short-day conditions (8-h light/16-h dark) at 22°C or long-day conditions (16-h light/8-h dark) at 25°C and 50-70% humidity. Seeds were treated in 10% (v/v) sodium hypochlorite, washed five times with sterile water, stratified at 4°C for 48 h, and plated on 0.5 x Murashige and Skoog (MS) medium with 0.8% (w/v) agar plates.

### 2.2 Promoter analysis

The genomic sequence was retrieved from the Ensembl Plants database based on the TAIR10 genome assembly (GCF_000001735.4) (Lamesch et al. 2012). The promoter region was defined as the 1104 bp sequence immediately upstream of the translation start site, corresponding to the genomic coordinates 2:7,707,955-7,709,058. *Cis*-regulatory elements were identified using PlantPAN 4.0 (Chow et al. 2024) applying a similarity score cutoff of 0.9 to ensure high confidence predictions. The identified binding sites were categorized by biological role by submitting the unique AGI codes of the predicted transcription factors to the TAIR GO annotation search to associate them with specific Gene Ontology terms. The functionally annotated promoter map was generated using Python (v3.11.13) with the pandas, Matplotlib, and DNA Features Viewer (Zulkower and Rosser 2020) libraries.

### 2.3 Seed germination, plant growth, morphological analysis and salt tolerance studies

For *in vitro* germination tests, surface-sterilized *A. thaliana* seeds, from wild-type (WT), *VLG*-knock-down (L1, L2, and L3) and *VLG*-overexpressing (OE1 and OE2) lines were plated in Petri dishes, in 0.5 x MS medium with 0.8% (w/v) agar, supplemented with 0, 50, 75, or 100 mM NaCl. The plates were incubated in a growth chamber under long-day conditions described above for 6 days. *A. thaliana* growth stages and parameters, such as germination, length of primary root and rosette and total exposed leaf area, was determined as described by Boyes et al. (2001). Germination was evaluated considering seedlings that reached Stage 1.0 (fully opened cotyledons) as germinated. Germination rates were then calculated as previously described by Song et al. (2020). For root elongation tests, seedlings were grown for 4-5 days on vertical plates with 0.5 x MS agar (supplemented with 1% sucrose), and transferred to MS agar plates with 0, 50, 75, or 100 mM NaCl. Images were captured on an Axiocam HRC CCD camera (Zeiss, http://www.zeiss.com/) using the Axiovision program (version 4.2), and the primary root length was then quantified using ImageJ software (Fiji v.2.15.0).

For salt tolerance tests of adult plants, WT, *VLG*-knock-down and *VLG*-overexpressing lines seeds were individually sown in 5-cm-diameter pots containing a 3:1:1 mix of soil, vermiculite and perlite. Salt stress was initiated four weeks after sowing, by the addition of 200 mM NaCl to irrigation water, and was maintained for three additional weeks. For biomass determination, the plants were removed from the soil and washed with deionized water. Plant biomass was first recorded as fresh weight (FW). Samples were then dried in an oven at 65°C for 72–96 h until a constant weight was reached, which was recorded as the dry weight (DW).

### 2.4 GUS staining

For *VLG* promoter activity observed by GUS staining tests, *proVLG*:*GUS* developing seedlings were processed as described by (Pagnussat et al. 2007). *ProVLG*:*GUS* seedlings were grown on 0.5 MS agar plates (supplemented with 1% sucrose), until Principal growth Stage 1.02 (Boyes et al. 2001), with two leaf primordia and then transferred to 0.5 x MS liquid medium with 0 or 100 mM NaCl. Pictures were taken after 30 and 120 min of treatment.

### 2.5 Measurement of membrane permeability

Membrane permeability was expressed as Relative Electrolyte leakage (Liu et al. 2012). Four-week-old WT, *VLG*-knock-down and *VLG*-overexpressing plants were treated with 0 or 200 mM NaCl for 3 weeks. Intact leaves were then excised from the plants. Five leaf discs, each 5 mm in diameter, were cut from the excised leaves using a cork borer and immediately floated in 1 ml of deionized water for 4 h, after which conductivity of the solution was measured in micro-ohms (M1) using a conductivity meter (Compact Conductivity Meter, LAQUAtwin-EC-11, HORIBA Scientific). Samples were then boiled for 20 min and cooled down to room temperature. The solution conductivity of the killed tissues (M2) was measured. Ion leakage was calculated as the ratio of M1 to M2. Three independent experiments were conducted with a minimum of three replicates.

### 2.6 Measurement of chloroplast pigment contents in *A. thaliana* leaves

For total chlorophyll extractions, four-week-old WT, *VLG*-knock-down and *VLG*-overexpressing plants were treated with 0 or 200 mM NaCl for 3 weeks. Approximately 0.5 g of leaves were soaked in 10 ml 80% (v/v) acetone for 48 h at 4°C in the dark. Absorbance values of the extracts were measured spectrophotometrically at 663, 646, and 470 nm, and relative chlorophyll contents were calculated according to Song et al. (2023):

Chlorophyll a = (12.21*OD_663_–2.81*OD_646_)*FW (g)

Chlorophyll b = (20.13*OD_646_–5.03*OD_663_)*FW (g)

Carotenoid = ((1000*OD_470_–3.27* Chlorophyll a-104* Chlorophyll b)/229)*FW (g)

To estimate the anthocyanin concentration, whole rosettes were incubated in 80% (v/v) methanol with 0.01 M HCl (0.45 g of tissue per 10 ml of solution) at 4°C in the dark for 24 h. Then, 300 μl of extract was added to 200 μl of H_2_O and 100 μl of chloroform, and centrifuged (13,000 g for 10 min). Anthocyanin content was estimated by calculating the difference between the absorbance values at 540 and 630 nm in the methanolic phase (Del Castello et al. 2021).

### 2.7 *In situ* determination of reactive oxygen species (ROS)

Detection of hydrogen peroxide (H_2_O_2_) and superoxide anion (O_2_^−^) in plant tissues was performed by 3,3′-diaminobenzidine (DAB) and nitroblue tetrazolium (NBT) staining, respectively, according to Villarreal et al. (2009), with minor modifications. Briefly, four-week-old WT, *VLG*-knock-down, and *VLG*-overexpressing plants were treated with 0 or 200 mM NaCl for 24 h. Leaves were incubated overnight in the dark in a solution containing 0.25 mg/ml of DAB in 50 mM sodium acetate buffer (pH 5.4), or for 2 h with 0.5 mg/ml of NBT in 25 mM HEPES buffer (pH 7.0) at 25°C. Afterwards, the solution was removed and tissues were decolorated with 90% (v/v) ethanol overnight at 25°C to remove chlorophyll. Images were taken with a binocular SMZ800 (Nikon) and a CoolSnap Pro camera (MediaCybernetics, TX). The DAB or NBT signal was digitally quantified as the average pixel light value using Image J software (Fiji v.2.15.0). Three independent experiments were conducted with 36 replicates in total per treatment.

### 2.8 Measurement of Na^+^ and K^+^ contents in *A. thaliana* leaves

Four-week-old WT, *VLG*-knock-down, and *VLG*-overexpressing plants were treated with 0 or 200 mM NaCl for 3 weeks. The leaves were harvested and dried for 72 h at 65°C, and then ground to powder. 200 mg of this powdered tissue was weighed into individual digestion tubes. Each sample received 2 ml concentrated perchloric acid (HClO_4_) and 2 ml concentrated nitric acid (HNO_3_), and were digested in a VelpScientific unit for 1 h at 120°C followed by 1 h at 150°C. Mineralized samples were diluted to 100 ml with deionized water. For K^+^ determination, a 0.5 ml aliquot was mixed with 0.5 ml lanthanum chloride (LaCl_3_) and 9.0 ml deionized water. For Na^+^, a 2.0 ml aliquot was combined with 8.0 ml potassium chloride (KCl). Analyses were performed by atomic emission spectrometry (PerkinElmer AA700) and concentrations were calculated with LCsolution software (Shimadzu Corp). A minimum of 30 plants were assessed in each treatment. Two independent experiments were conducted with three replicates per treatment.

### 2.10 RNA isolation and qRT-PCR analysis

Total RNA was isolated from four-week-old *A. thaliana* leaves with or without 200 mM NaCl treatment for 3 days; with TRIzol reagent (Invitrogen, USA). The quantity and quality of RNA were assessed by spectrophotometry (OD_260/280_) and agarose gel electrophoresis. cDNA was synthesized from 1 μg of DNase I-treated total RNA (Promega) using ImProm-II Reverse Transcriptase (Promega) and random hexamer primers. qPCR reactions were performed using FastStart Universal SYBR Green Master (ROX) PCR mix (Roche) in a StepOne machine (Applied Biosystems). The amplification conditions were 95 °C for 10 min, followed by 40 cycles of amplification (95 °C for 15 s, 60 °C for 30 s) and reactions were conducted in triplicates. The relative expression of genes was calculated using the 2^ΔΔCt^ method (Livak and Schmittgen 2001). Data are presented as relative expression levels normalized to the reference gene *Actin 2*, based on three independent experiments. The primers used for qRT-PCR are shown in Supplemental Table S1.

### 2.11 Statistical analysis

All experiments were independently repeated at least three times, and each assay was performed in triplicate. The resulting data were analyzed statistically using GraphPad Prism version 6.01. Statistical differences among groups were analyzed by ANOVA followed by Tukey’s or Dunnett’s post hoc tests, with P < 0.05 considered significant.

## 3 Results

### 3.1 VLG is transcriptionally activated under salt stress conditions

Given the proposed involvement of several DC1 domain proteins in abiotic stress responses, we first examined whether VLG expression is modulated by salinity. *VLG* transcript levels increased significantly in *A. thaliana* seedlings upon NaCl treatment (Figure 1A). Consistent with this observation, analysis of *VLG* promoter activity revealed that *GUS* expression was strongly induced in *proVLG:GUS* seedlings treated with 100 mM NaCl (Figure 1B), supporting transcriptional activation of *VLG* under salt stress conditions.

**FIGURE 1.**
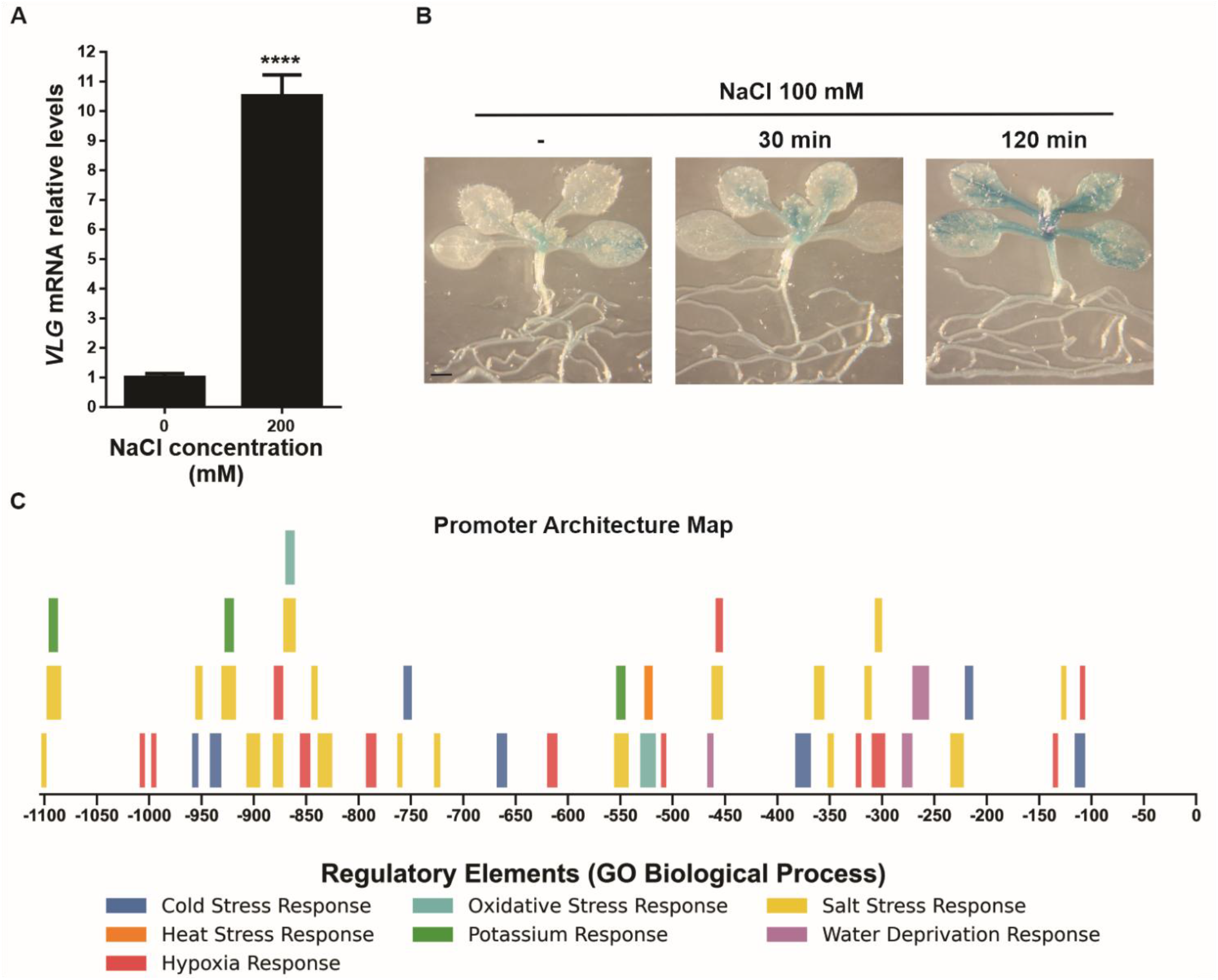
*VLG* is transcriptionally induced by salt stress, and its promoter contains multiple stress-responsive cis-elements. Four-week-old WT plants were treated with or without 200 mM NaCl for 3 days. (A) The expression levels of *VLG* were measured by qRT-PCR. Unpaired t test, p<0.0001 (****). (B) The activity of the *VLG* promoter in *proVLG:GUS* seedlings treated with or without 100 mM NaCl for 30 or 120 minutes. Three independent experiments were done with similar results. Scale bar = 1 mm. (C) *In silico* analysis of cis-acting elements of *VLG* gene promoter. The line represents the upstream 1104 bp promoter of the *VLG* gene. Boxes indicate the transcription factors (TFs) predicted to bind to specific cis-elements, colored according to the Gene Ontology functional categories.

To further explore the regulatory basis of this response, we analyzed the VLG promoter sequence *in silico*. Multiple *cis*-acting elements associated with abiotic stress signaling were identified, including motifs related to osmotics stress, oxidative stress, heat stress, and drought responses (Figure 1C, Supplementary Table S2). Notably, the promoter contains several regulatory elements previously associated with salinity-responsive transcriptional networks, including W-boxes, NAC-binding sites, and bHLH-related motifs (Balazadeh et al. 2010, Liu et al. 2014, Ma and Hu 2024). Together, these results indicate that VLG is transcriptionally responsive to salinity and support its potential involvement in plant adaptive responses to salt stress.

### 3.2 VLG levels regulates salt tolerance throughout plant development

The salt-induced expression of *VLG* and the presence of stress-responsive elements in its promoter prompted us to investigate whether VLG plays a functional role in plant tolerance to salinity.

To assess this, we first analyzed seed germination in WT, *VLG*-knock-down (L1–L3) and *VLG*-overexpressing (OE1, OE2) *A. thaliana* lines under increasing NaCl concentrations (0, 50, 75 and 100 mM; Figure 2A, B). Without NaCl addition, all the lines showed similar germination rates (Figure 2A, B). Addition of salt resulted in lower germination rates for all the lines, however, *VLG*-knock-down lines showed significant lower values, compared with WT plants. On the other hand, germination rates for *VLG*-overexpressing lines were similar to WT plants (Figure 2A, B). We next analyzed primary root growth under the same salt stress conditions (Figure 2C, D). Under normal growth conditions, all genotypes were indistinguishable (Figure 2C, D). However, upon exposure to salt stress, *VLG*-knock-down lines exhibited significantly reduced primary root elongation compared with both WT and *VLG*-overexpressing plants (Figure 2C, D).

**FIGURE 2.**
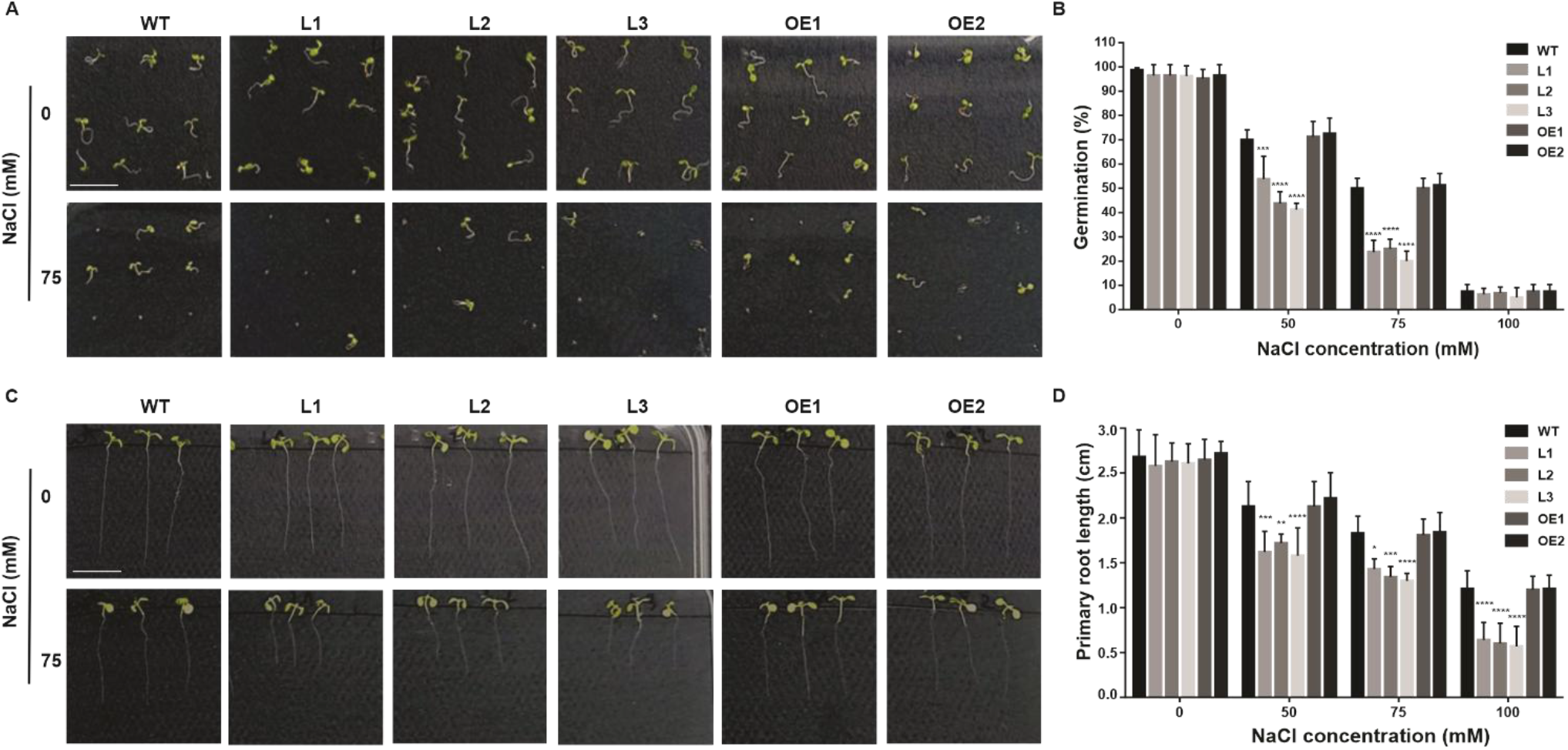
Altered *VLG* expression affects seed germination and primary root growth under salt stress. (A) Representative phenotypes and (B) germination rates of WT, *VLG*-knock-down (L1, L2, L3) and *VLG*-overexpressing (OE1, OE2) seedlings grown on MS medium supplemented with 0, 50, 75 and 100 mM NaCl for 6 days. (C) Representative phenotypes and (D) measurement of primary root length of WT, *VLG*-knock-down and *VLG*-overexpressing seedlings grown on MS medium supplemented with 0, 50, 75 and 100 mM NaCl for 10 days. ANOVA and Dunnett’s multiple comparisons test, p<0.05 (*), p<0.01 (**), p<0.001 (***), p<0.0001 (****). All experiments were repeated at least three times. Scale bar = 1 cm.

Next, we analyzed the performance of four-week-old plants under control or salt stress conditions. WT, *VLG*-knock-down, and *VLG*-overexpressing plants were grown in soil for four weeks under short-day (Figure 3) or long-day (Figure S1) photoperiods and subsequently irrigated with 0 or 200 mM NaCl for an additional three weeks. Under control conditions, *VLG*-knock-down plants showed a mild epinastic phenotype, consistent with previous reports (Amigo et al., 2025), although no significant differences in rosette area were observed among genotypes at either 4 or 7 weeks of growth (Figure 3A, B, Figure S1). However, under salt stress conditions, *VLG*-knock-down plants exhibited a salt-sensitive phenotype, characterized by a smaller rosette area, fewer leaves, reduced height and lower fresh and dry weight than WT plants under both photoperiods tested (Figure 3, Figure S1). Surprisingly, *VLG*-overexpressing plants displayed a salt stress resistant phenotype, with a larger rosette area, more leaves, taller height and higher fresh and dry weight than WT plants (Figure 3, Figure S1). Although salinity reduced biomass accumulation in all genotypes, this reduction was substantially more severe in *VLG*-knock-down lines.

**FIGURE 3.**
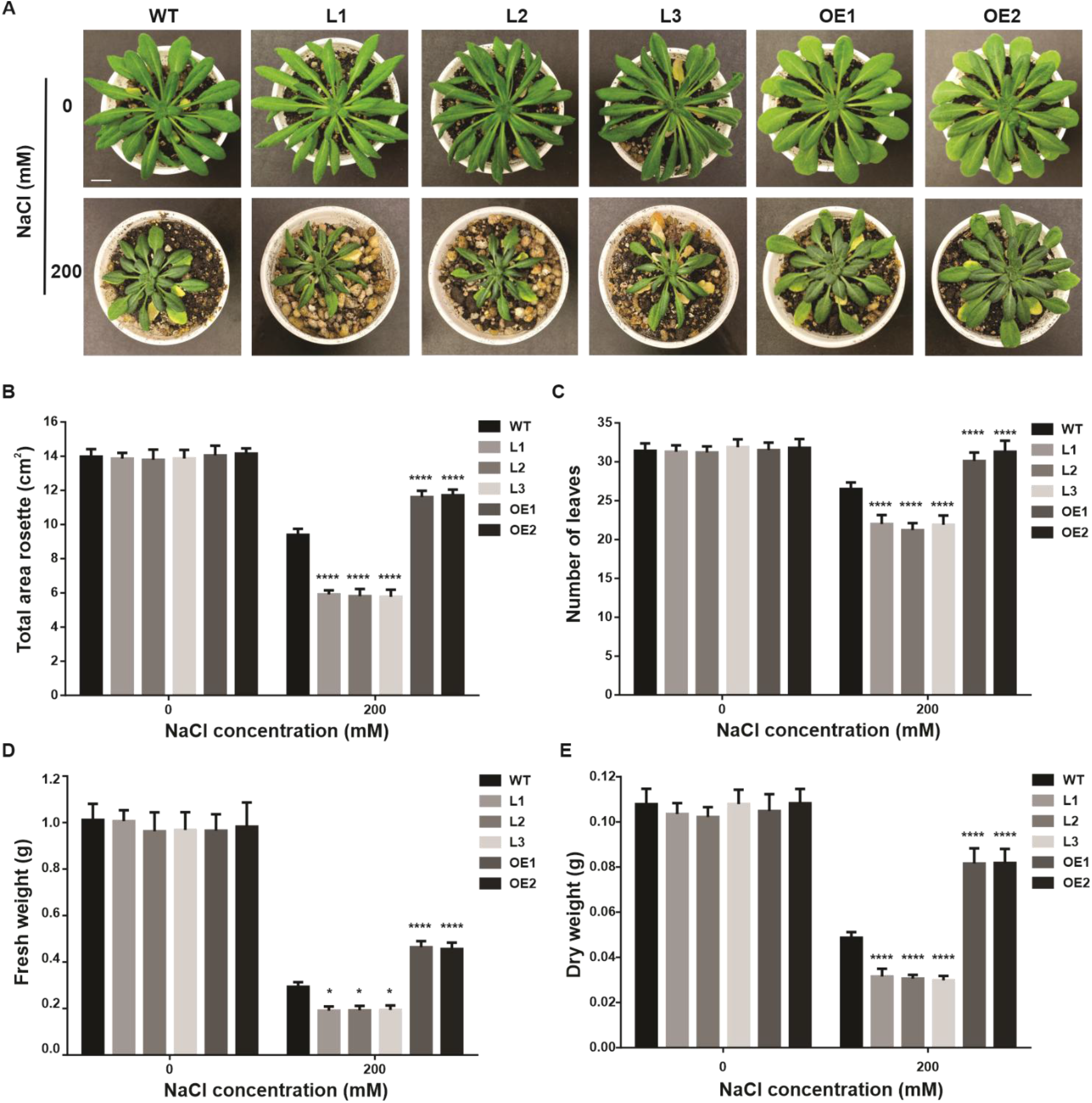
Altered *VLG* expression affects plant growth and biomass accumulation under salt stress. (A) Representative phenotypes, (B) rosette area, (C) leaf number, (D) fresh weight, and (E) dry weight of four-week-old WT, *VLG*-knock-down (L1, L2, L3) and *VLG*-overexpressing (OE1, OE2) plants grown under short-day condition with or without 200 mM NaCl treatment for 21 days. ANOVA and Dunnett’s multiple comparisons test, p<0.05 (*), p<0.0001 (****). Three independent experiments were done with similar results. Scale bar = 1 cm.

Taken together, these findings indicate that VLG expression influences seed germination, primary root elongation, and overall plant growth, supporting an active role for VLG in salinity tolerance in *A. thaliana*.

### 3.3 VLG maintains photosynthetic and ionic homeostasis under saline conditions

To investigate the physiological basis underlying the altered salinity responses observed in *VLG* transgenic lines, we examined photosynthetic pigment accumulation and ion homeostasis in WT, VLG-knock-down, and VLG-overexpressing plants exposed to salt stress. Four-week-old WT, *VLG*-knock-down and *VLG*-overexpressing plants grown under short-day (Figure 4) or long-day (Figure S2) photoperiod conditions were treated for 3 weeks with 200 mM NaCl. Salt stress caused a reduction of chloroplast pigments across all lines (Figure 4A-C), however, chlorophyll a and carotenoids were significantly lower in *VLG*-knock-down plants compared to WT, whereas *VLG*-overexpressing plants maintained pigment levels comparable to WT. No differences in the levels of chloroplast pigments were observed between the different lines under normal growth conditions (Figure 4A-C). On the other hand, anthocyanin levels were significantly higher in *VLG*-knock-down lines under normal conditions (Figure 4D). Upon salt stress, anthocyanin levels increased in all the lines, remaining significantly higher in *VLG*-knock-down plants relative to WT (Figure 4D). Similar results were observed under long-day conditions in all studied lines (Figure S2). These results indicate that reduced VLG expression is associated with enhanced physiological stress and impaired maintenance of photosynthetic integrity under salinity.

**FIGURE 4.**
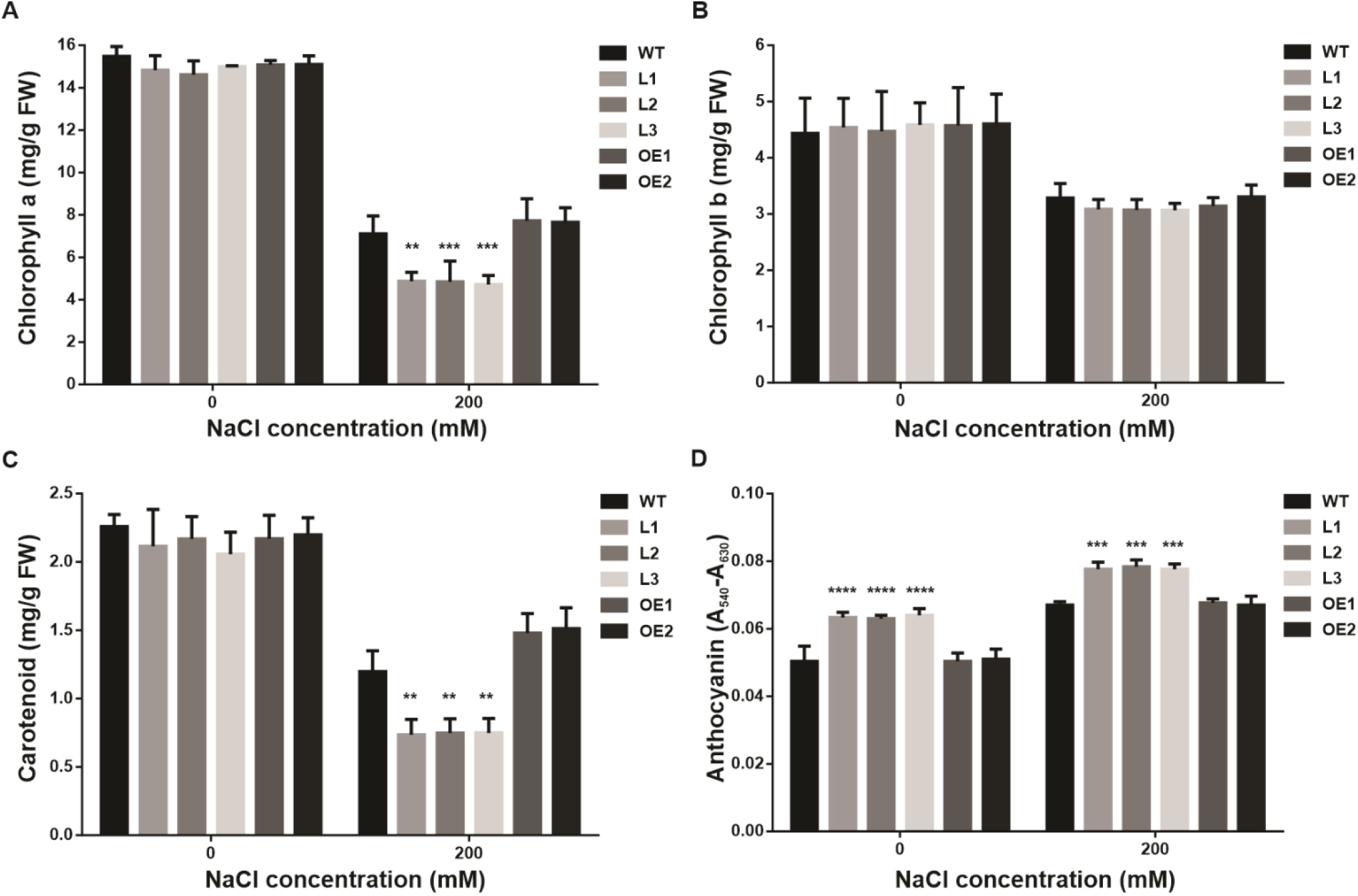
Altered *VLG* expression affects photosynthetic pigments and anthocyanin accumulation under salt stress. (A) The contents of chlorophyll a, (B) chlorophyll b, (C) carotenoid, and (D) anthocyanin in leaves of four-week-old WT, *VLG*-knock-down (L1, L2, L3) and *VLG*-overexpressing (OE1, OE2) plants grown under short-day condition with or without 200 mM NaCl treatment for 21 days. ANOVA and Dunnett’s multiple comparisons test, p<0.01 (**), p<0.001 (***), p<0.0001 (****). Three independent experiments were done with similar results.

We next analyzed Na^+^ and K^+^ accumulation to determine whether VLG affects ionic homeostasis during salt stress (Figure 5). Under control conditions, no significant differences in ion content were observed between transgenic and WT plants. Saline conditions increased Na^+^ accumulation and reduced K^+^ levels in all genotypes; however, the magnitude of these changes differed markedly depending on *VLG* expression levels. WT plants accumulated the highest levels of Na^+^ together with the strongest reduction in K^+^ content, resulting in elevated the Na^+^/K^+^ ratio. In contrast, *VLG*-overexpressing plants maintained significantly lower Na^+^ levels and higher K^+^ concentrations, leading to substantially lower Na^+^/K^+^ ratios relative to WT plants. *VLG*-knock-down lines displayed intermediate values, although they accumulated less Na^+^ than WT, their Na^+^/K^+^ ratios remained higher than those of the overexpressing plants.

**FIGURE 5.**
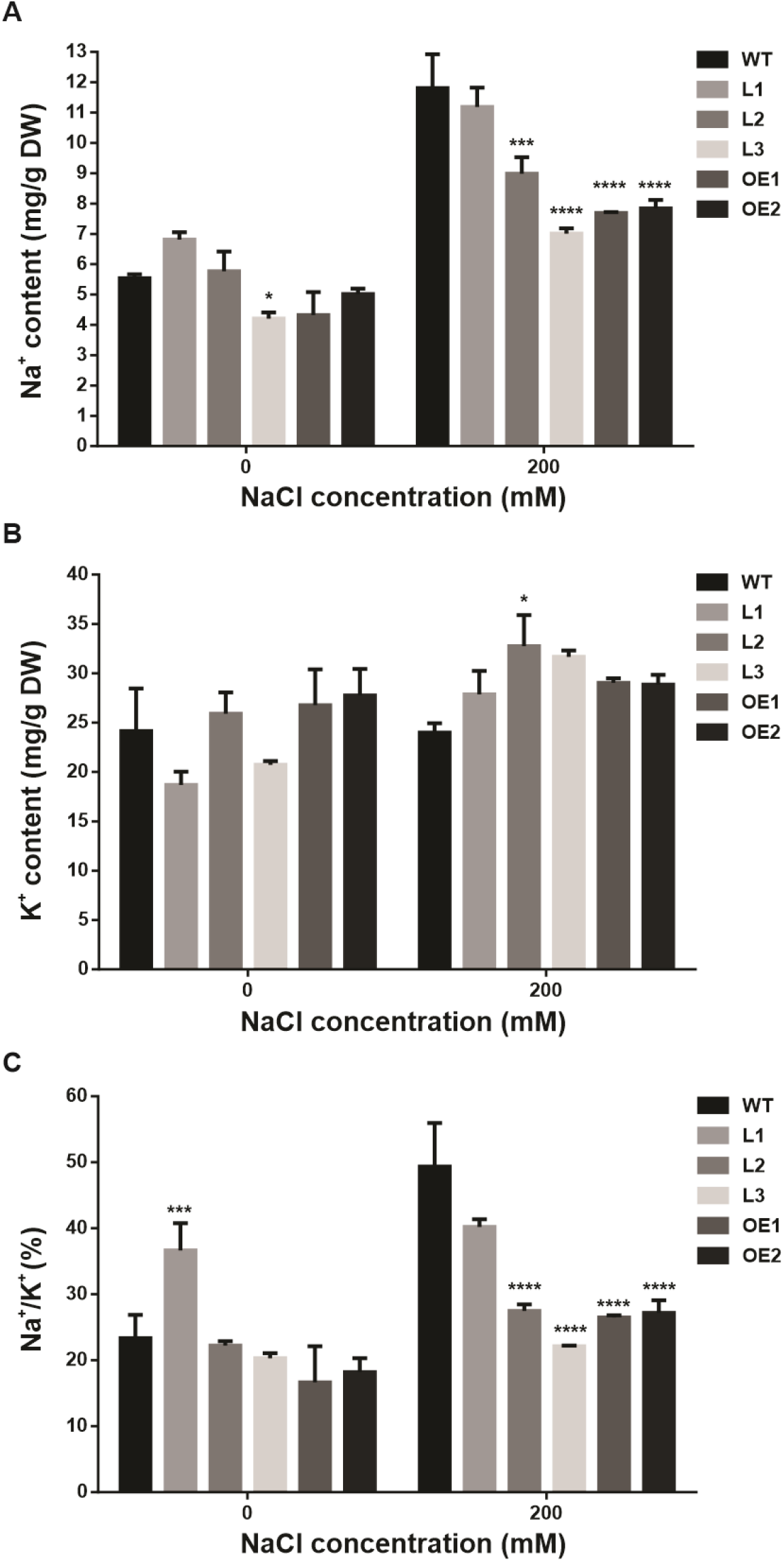
Altered *VLG* expression affects Na^+^ and K^+^ content under saline conditions. (A) Na^+^ content, (B) K^+^ content, and (C) Na^+^/K^+^ ratio in leaves of four-week-old WT, *VLG*-knock-down (L1, L2, L3) and *VLG*-overexpressing (OE1, OE2) plants grown under short-day condition with or without 200 mM NaCl treatment for 21 days. ANOVA and Dunnett’s multiple comparisons test, p<0.05 (*), p<0.01 (**), p<0.001 (***), p<0.0001 (****). Two independent experiments were done with similar results.

Notably, despite accumulating less Na^+^ than WT plants, *VLG*-knock-down lines exhibited the strongest growth inhibition and physiological impairment under salinity. This observation suggests that the hypersensitive phenotype associated with reduced VLG expression cannot be attributed solely to Na^+^ accumulation, pointing instead to a broader role of VLG in stress adaptation and cellular homeostasis. Together with the altered photosynthetic pigment composition and enhanced anthocyanin accumulation observed in *VLG*-knock-down plants, these results indicate that VLG contributes to the maintenance of physiological integrity under saline conditions.

### 3.4 VLG prevents ROS accumulation and reduces salt-induced membrane damage

Because oxidative stress is a major component of salt-induced cellular damage, we then analyzed ROS accumulation and membrane integrity in *VLG* transgenic lines exposed to salinity.

DAB and NBT staining revealed increased accumulation of H_2_O_2_ and O_2_^−^, respectively, in *VLG*-knock-down plants after 24 h of salt treatment (Figure 6A–D). In contrast, *VLG*-overexpressing plants displayed ROS levels comparable to WT plants under the same conditions. These results indicate that reduced *VLG* expression enhances oxidative imbalance during salt stress.

**FIGURE 6.**
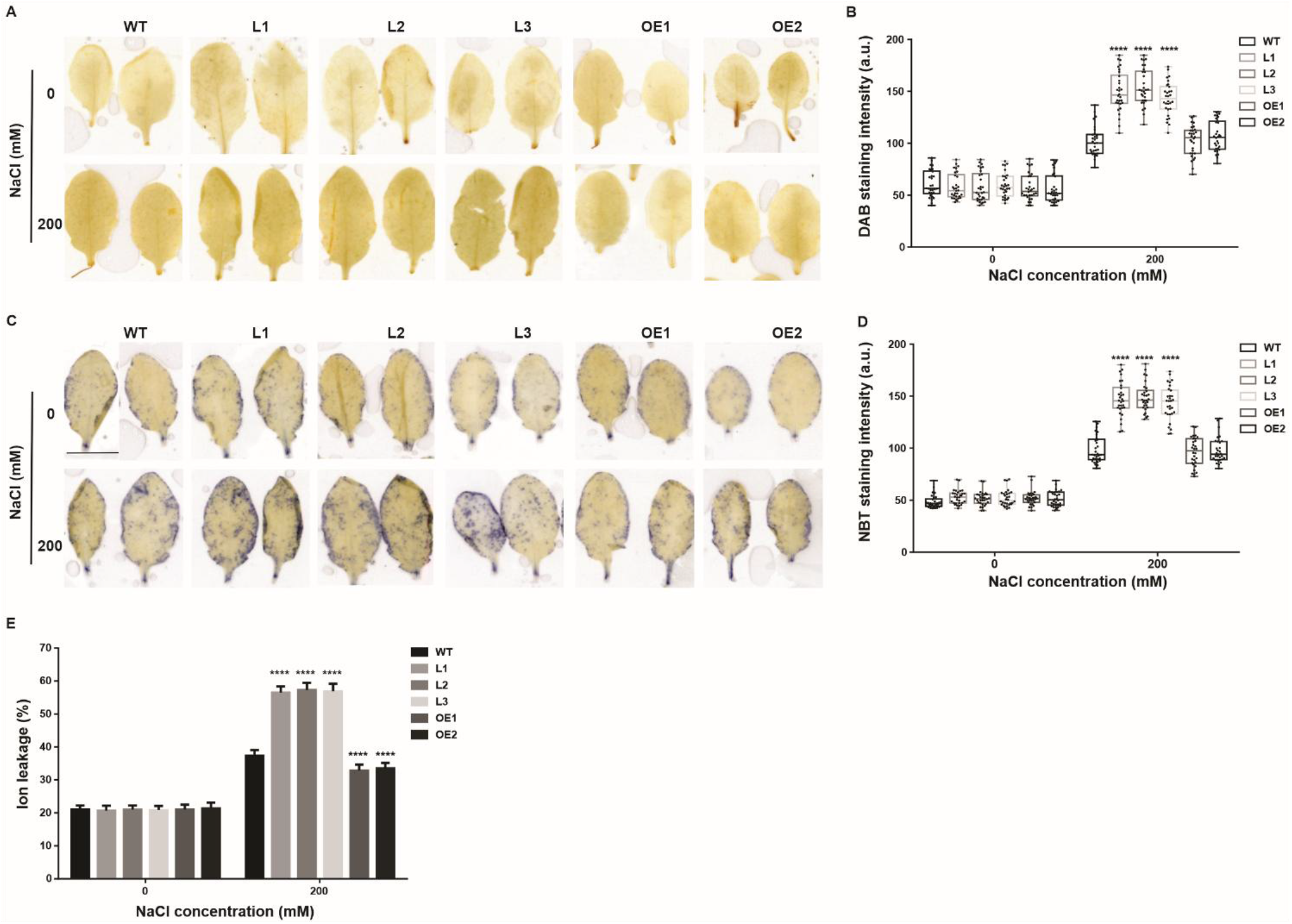
Altered *VLG* expression affects ROS accumulation and membrane integrity under salt stress. (A) Representative DAB staining for H_2_O_2_, accumulation and (C) NBT staining for O^-^ accumulation, together with their corresponding quantifications (B, D) in leaves of four-week-old WT, *VLG*-knock-down (L1, L2, L3) and *VLG*-overexpressing (OE1, OE2) plants grown under short-day condition with or without 200 mM NaCl treatment for 24hs. or for 3 weeks. (E) Ion leakage measurements in leaves of WT, *VLG*-knock-down, and *VLG*-overexpressing plants treated with or without 200 mM NaCl for 3 weeks under the same growth conditions. ANOVA and Dunnett’s multiple comparisons test, p<0.0001 (****). Three independent experiments were done with similar results. Scale bar = 1 cm.

To further assess cellular damage under salinity, membrane integrity was evaluated through electrolyte leakage measurements. Salt treatment caused a marked increase in ion leakage in *VLG*-knock-down plants relative to WT plants, whereas *VLG*-overexpressing lines displayed lower electrolyte leakage than WT plants under saline conditions (Figure 6E; Figure S3). Together, these findings show that reduced *VLG* expression enhances salt-induced oxidative damage and loss of membrane integrity.

### 3.5 VLG modulates the expression of key salt-responsive genes

Given the physiological alterations observed in *VLG* transgenic lines under salinity, we finally explored whether *VLG* expression levels affect the transcriptional response to salt stress. As expected, *VLG* transcript levels increased significantly in four-week-old WT plants after exposure to 200 mM NaCl for 3 days, whereas *VLG*-overexpressing lines displayed an even stronger induction under the same conditions (Figure S4). We then analyzed the expression of key salt stress–related genes associated with stress signaling and ionic homeostasis by qRT-PCR. Four-week-old plants were treated with or without 200 mM NaCl for 3 days, and the expression levels of key regulators of ionic homeostasis -*Rap2*.*6, Rap2*.*6L, LTL1, SOS1, SOS2, NHX1, NHX3, NHX5*, and *HKT1-* were measured (Figure 7). Under control conditions, no significant differences in transcript levels were detected among genotypes.

**FIGURE 7.**
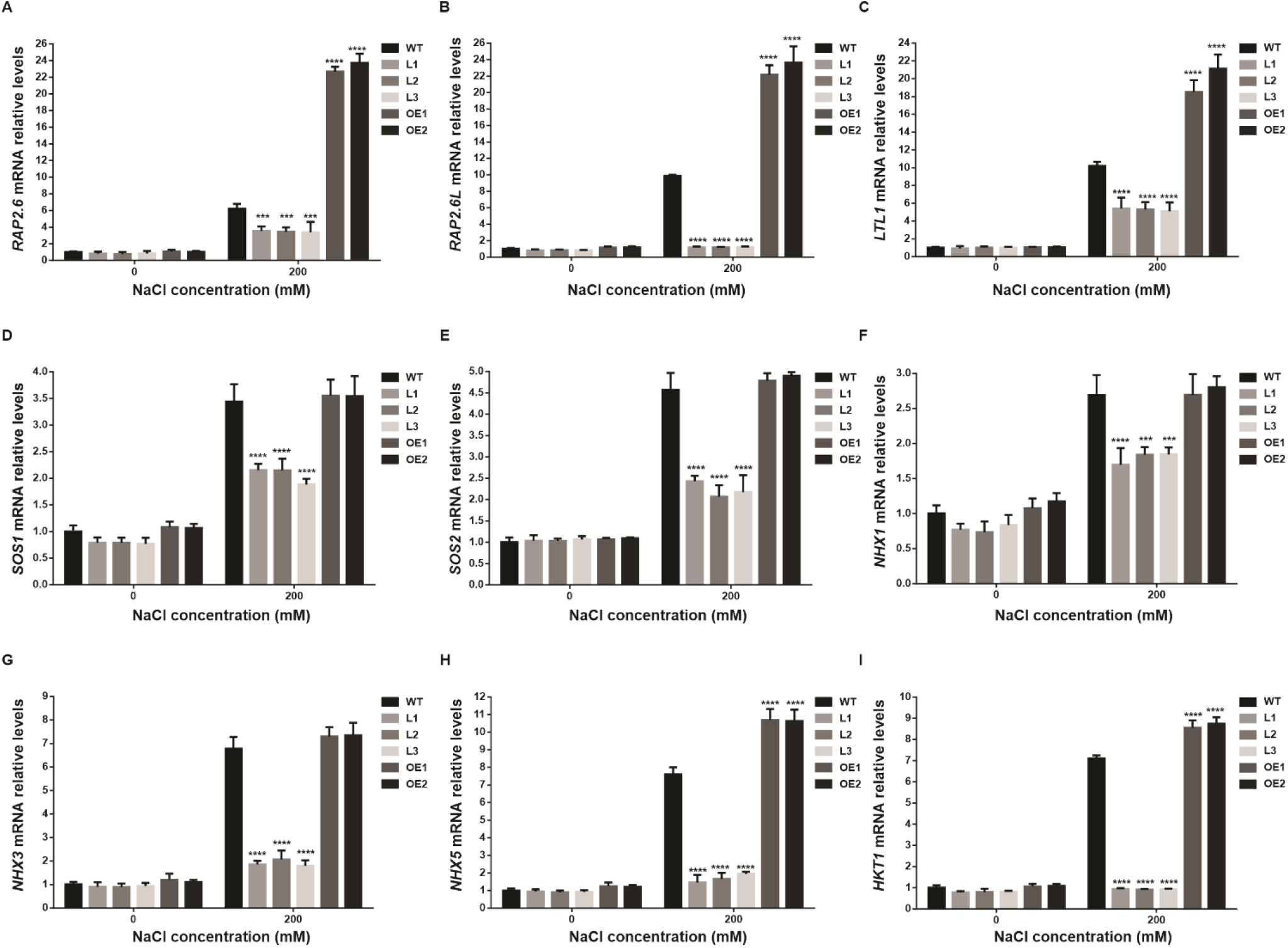
Altered *VLG* expression modulates the transcriptional response to salinity. Four-week-old WT, *VLG*-knock-down (L1, L2, L3) and *VLG*-overexpressing plants (OE1, OE2) were treated with or without 200 mM NaCl treatment for 3 days. The expression levels of (A) *RAP2*.*6*, (B) *RAP2*.*6L*, (C) *LTL1*, (D) *SOS1*, (E) *SOS2*, (F) *NHX1*, (G) *NHX3*, (H) *NHX5*, and (I) *HKT1* were measured by qRT-PCR. ANOVA and Dunnett’s multiple comparisons test, p<0.05 (*), p<0.01 (**), p<0.001 (***), p<0.0001 (****). Three independent experiments were done with similar results.

Salt treatment induced the expression of all marker genes in WT, *VLG*-knock-down, and *VLG*-overexpressing plants; however, the magnitude of this induction differed among genotypes (Figure 7). Compared with WT plants, *VLG*-knock-down lines exhibited reduced expression of several salt-responsive genes under saline conditions. In contrast, *Rap2*.*6, Rap2*.*6L, LTL1, NHX5*, and *HKT1* showed significantly higher transcript levels in *VLG*-overexpressing plants relative to WT plants following salt treatment (Figure 7A, B, C, H, I). Together, these results indicate that *VLG* expression levels influence the transcriptional response to salinity, particularly affecting genes associated with stress signaling and ionic homeostasis.

## 4 Discussion

Salinity is an increasingly relevant environmental constraint affecting agricultural productivity worldwide, particularly in arid and semi-arid regions where progressive soil salinization limits the availability of cultivable land (Yang and Guo 2018, Van Zelm et al. 2020). Plant adaptation to saline environments requires the coordination of multiple physiological and cellular processes, including ion homeostasis, oxidative stress control, membrane function, and stress-responsive transcriptional programs. Growing evidence indicates that DC1 domain proteins contribute to plant adaptation to saline environments across multiple plant species. However, the physiological and molecular functions of most DC1 proteins remain largely unexplored (Li et al. 2010, Gao et al. 2016, Li et al. 2023, Kang et al. 2024, Du et al. 2026).

Among DC1 proteins, VLG represents a particularly interesting member of the family. VLG was previously characterized as essential for gametophyte development and central vacuole formation in *A. thaliana* (D’Ippólito et al. 2017), but the present work reveals a broader physiological role for this protein under saline conditions. Altered *VLG* expression strongly affected plant performance under saline conditions throughout development, from seed germination to adult growth. Reduced *VLG* expression increased salt sensitivity, whereas *VLG* overexpression improved growth and biomass accumulation under salinity, supporting a broader role for this DC1 domain protein beyond reproductive development.

One of the most evident consequences of altered *VLG* expression was the disruption of physiological homeostasis under salinity. *VLG*-knock-down plants displayed enhanced ROS accumulation, increased membrane damage, altered pigment composition, and impaired growth under saline conditions, whereas *VLG*-overexpressing plants maintained lower oxidative damage and improved growth performance. The increased accumulation of H_2_O_2_ and O_2_^−^ observed in *VLG*-knock-down plants is particularly relevant considering the central role of ROS during salt stress. Although ROS participate in stress signaling and acclimation responses, excessive ROS accumulation promotes membrane lipid peroxidation, protein oxidation, and cellular damage, ultimately compromising photosynthetic performance and plant growth (Apel and Hirt 2004, Miller et al. 2010). Consistent with this, the enhanced electrolyte leakage and anthocyanin accumulation observed in *VLG*-knock-down plants suggest that reduced *VLG* expression compromises the ability to maintain redox homeostasis under saline conditions.

The ionic profiles observed in *VLG* transgenic lines further support this idea, although they also reveal a more complex physiological scenario. *VLG*-overexpressing plants maintained lower Na^+^/K^+^ ratios under salinity, consistent with improved ionic homeostasis. However, despite exhibiting stronger growth inhibition and oxidative damage, *VLG*-knock-down plants did not accumulate more Na^+^ than WT plants. This suggests that the hypersensitive phenotype associated with reduced *VLG* expression cannot be explained exclusively by altered Na^+^ accumulation. Instead, *VLG*-dependent salinity tolerance appears to involve the integration of multiple adaptive processes, including ionic balance, oxidative stress control, and stress-responsive transcriptional responses.

Consistent with the physiological and ionic phenotypes observed in *VLG* transgenic lines, several genes associated with salinity responses displayed altered expression patterns under salt stress. These included key regulators of ionic homeostasis such as *SOS1* and *SOS2*, components of the SOS pathway involved in Na^+^ extrusion (Yang and Guo 2018), as well as *HKT1*, which contributes to Na^+^ retrieval from the shoot and maintenance of Na^+^/K^+^ balance under saline conditions (Jaime-Pérez et al. 2017). In addition, the vacuolar and endosomal antiporters *NHX1, NHX3*, and *NHX5*, which participate in intracellular ion compartmentalization and pH regulation required for endomembrane homeostasis during stress (Bassil et al. 2011, Reguera et al. 2015), were also differentially regulated in response to altered *VLG* expression. Notably, stress-responsive transcription factors of the AP2/ERF family, such as *RAP2*.*6* and *RAP2*.*6L*, together with *LTL1*, a lipase previously associated with salt tolerance in yeast and plants (Naranjo et al. 2006, Krishnaswamy et al. 2011), showed reduced induction in *VLG*-knock-down plants and enhanced expression in *VLG*-overexpressing lines. These transcriptional profiles are consistent with the physiological behavior of the transgenic lines and further support the idea that VLG influences the capacity of plants to sustain coordinated acclimation responses under saline conditions.

This integrated behavior becomes particularly relevant considering the previously described features of VLG. VLG localizes to the endomembrane system and interacts with proteins associated with membrane trafficking, lipid metabolism, and transcriptional regulation (D’Ippólito et al. 2017). In this context, the altered expression of genes such as *NHX5* and *HKT1*, together with the oxidative and physiological phenotypes observed in *VLG* transgenic lines, points to a role for VLG in processes linked to endomembrane homeostasis and stress-associated cellular plasticity under saline conditions. This interpretation is consistent with the proposed scaffold-like behavior of several DC1 proteins and may explain the broad physiological effects associated with altered *VLG* expression. Rather than acting exclusively through Na^+^ exclusion mechanisms, *VLG* may contribute to the coordination of multiple cellular responses required to sustain physiological performance during salinity stress. The broad expression pattern of *VLG* in sporophytic tissues, together with its effects on growth, oxidative balance, and stress-responsive gene expression, further supports the idea that this protein participates in processes extending beyond gametophyte development. In this context, the present work expands the functional scope of DC1 domain proteins and highlights VLG as a candidate component linking endomembrane-associated processes with physiological adaptation to salinity stress in *A. thaliana*.

## Acknowledgments

We would like to thank Dr. Ayelen Distéfano for her assistance with the ion leakage assay, and Sergio Batista and Mariano Martinez for greenhouse assistance.

## Author contributions

**Natalia Amigo**: writing - original draft, investigation, formal analysis, data curation. **Fernanda Marchetti**: methodology, investigation. **Salvador Lorenzani**: methodology, investigation, validation. **Leonardo A. Arias**: methodology, investigation, **Juan Ignacio Poo**: methodology, investigation. **Milagros Escoriza**: methodology, investigation. **María Elisa Picco:** methodology, investigation. **María Cecilia Terrile**: writing - review & editing, supervision. **Diego F. Fiol**: writing - review & editing, supervision, resources, project administration, funding acquisition. All authors read and approved the manuscript.

## Generative AI statement

The authors used generative AI tools (e.g., ChatGPT, Perplexity) solely for language polishing. All original content and scientific contributions are exclusively those of the authors. The authors reviewed and approved all revisions and accept full responsibility for the manuscript.

## Data Availability Statement

The data used to support the findings of this study are available from the corresponding author upon request.

## Supporting Information

Additional supporting information can be found online in the Supporting Information section. **Supplementary FIGURE S1**. Altered *VLG* expression affects plant growth under salt stress during long-day conditions. (A) Representative phenotypes, (B) plant height, (C) fresh weight, and (D) dry weight of four-week-old WT, *VLG*-knock-down and *VLG-*overexpressing plants grown under long-day condition with or without 200 mM NaCl treatment for 21 days. ANOVA and Dunnett’s multiple comparisons test, p<0.05 (*), p<0.01 (**), p<0.0001 (****). Three independent experiments were done with similar results. Scale bar = 5 cm.

**Supplementary FIGURE S2**. Altered *VLG* expression affects photosynthetic pigment and anthocyanin accumulation under salt stress during long-day conditions. (A) Representative phenotypes, (B) chlorophyll a, (C) chlorophyll b, (D) carotenoid, and (E) and anthocyanin contents in leaves of four-week-old WT, *VLG*-knock-down (L1, L2, L3) and *VLG*-overexpressing (OE1, OE2) plants grown under long-day condition, with or without 200 mM NaCl treatment for 21 days. ANOVA and Dunnett’s multiple comparisons test, p<0.05 (*), p<0.0001 (****). Three independent experiments were done with similar results. Scale bar = 1 cm.

**Supplementary FIGURE S3**. Altered *VLG* expression affects membrane integrity under salt stress during long-day conditions. Ion leakage in WT, *VLG*-knock-down and *VLG*-overexpressing plants with or without salt treatment. Ion leakage was measured in leaves of four-week-old WT, *VLG*-knock-down (L1, L2, L3) and *VLG*-overexpressing (OE1, OE2) plants grown under long-day condition with or without 200 mM NaCl treatment for 3 weeks. ANOVA and Dunnett’s multiple comparisons test p<0.01 (**), p<0.0001 (****). Three independent experiments were done with similar results.

**Supplementary FIGURE S4**. Salt stress induces *VLG* expression in WT and *VLG*-overexpressing plants. Four-week-old WT, *VLG*-knock-down (L1, L2, L3), and *VLG*-overexpressing (OE1, OE2) plants were treated with or without 200 mM NaCl treatment for 3 days. The expression levels of *VLG* were measured by qRT-PCR. ANOVA and Dunnett’s multiple comparisons test, p<0.05 (*), p<0.001 (***), p<0.0001 (****). Three independent experiments were done with similar results.

**Supplementary TABLE S1**. Primers used for q-PCR.

**Supplementary TABLE S2**. Transcription factors identified in the promoter region of *VLG*.

