## Supplementary Figs S1-S4 and Tables S1, S2 for "The DC1 domain protein Vacuoleless Gametophytes positively regulates salt stress tolerance in *Arabidopsis thaliana*"

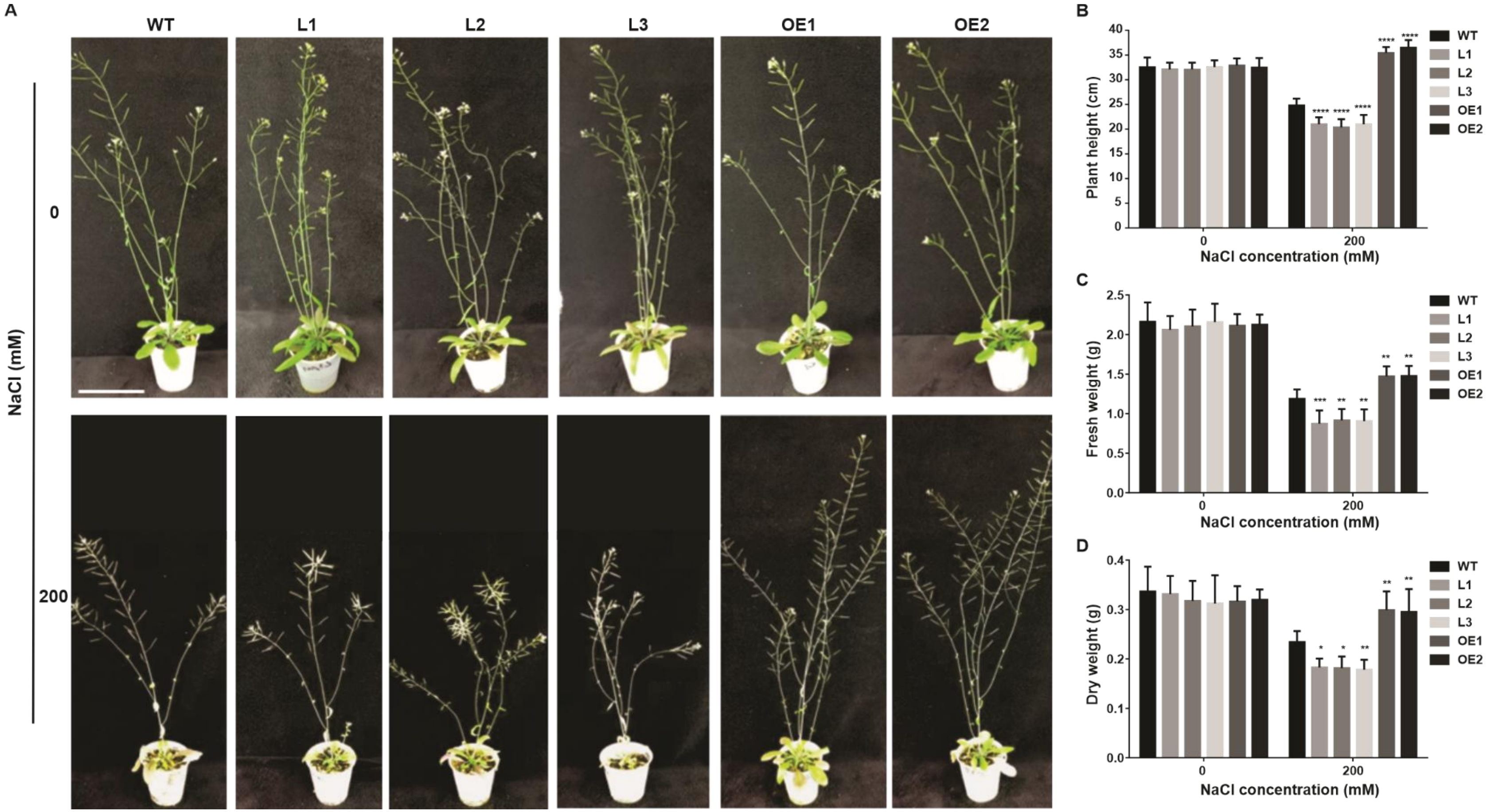


**Supplementary FIGURE S1.** Altered *VLG* expression affects plant growth under salt stress during long-day conditions. (A) Representative phenotypes, (B) plant height, (C) fresh weight, and (D) dry weight of four-week-old WT, *VLG*-knock-down and *VLG*-overexpressing plants grown under long-day condition with or without 200 mM NaCl treatment for 21 days. ANOVA and Dunnett's multiple comparisons test, $p<0.05$ (*), $p<0.01$ (**), $p<0.0001$ (****). Three independent experiments were done with similar results. Scale bar = 5 cm.

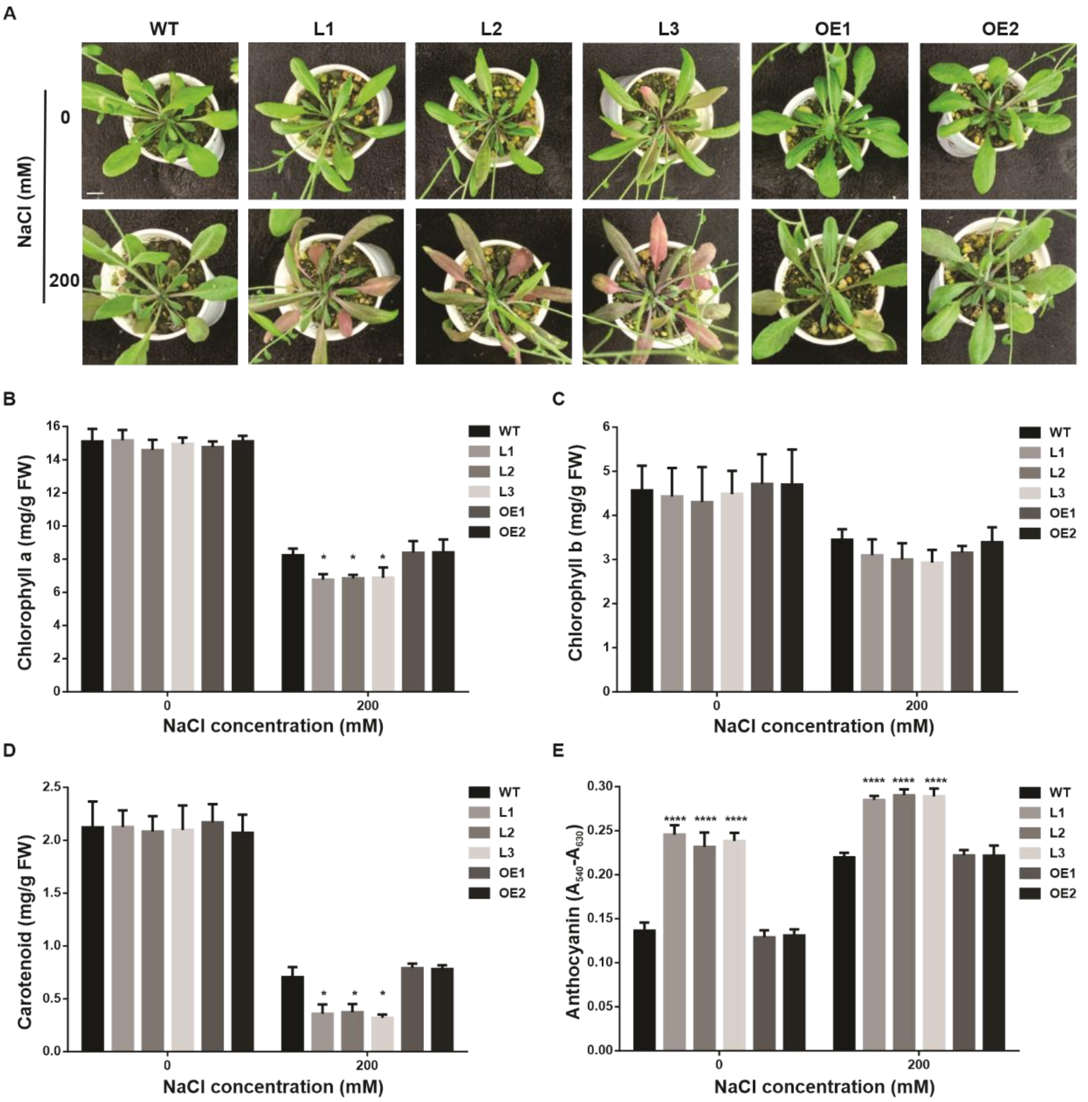


**Supplementary FIGURE S2.** Altered *VLG* expression affects photosynthetic pigment and anthocyanin accumulation under salt stress during long-day conditions. (A) Representative phenotypes, (B) chlorophyll a, (C) chlorophyll b, (D) carotenoid, and (E) and anthocyanin contents in leaves of four-week-old WT, *VLG*-knock-down (L1, L2, L3) and *VLG*-overexpressing (OE1, OE2) plants grown under long-day condition, with or without 200 mM NaCl treatment for 21 days. ANOVA and Dunnett's multiple comparisons test, p<0.05 (*), p<0.0001 (****). Three independent experiments were done with similar results. Scale bar = 1 cm.

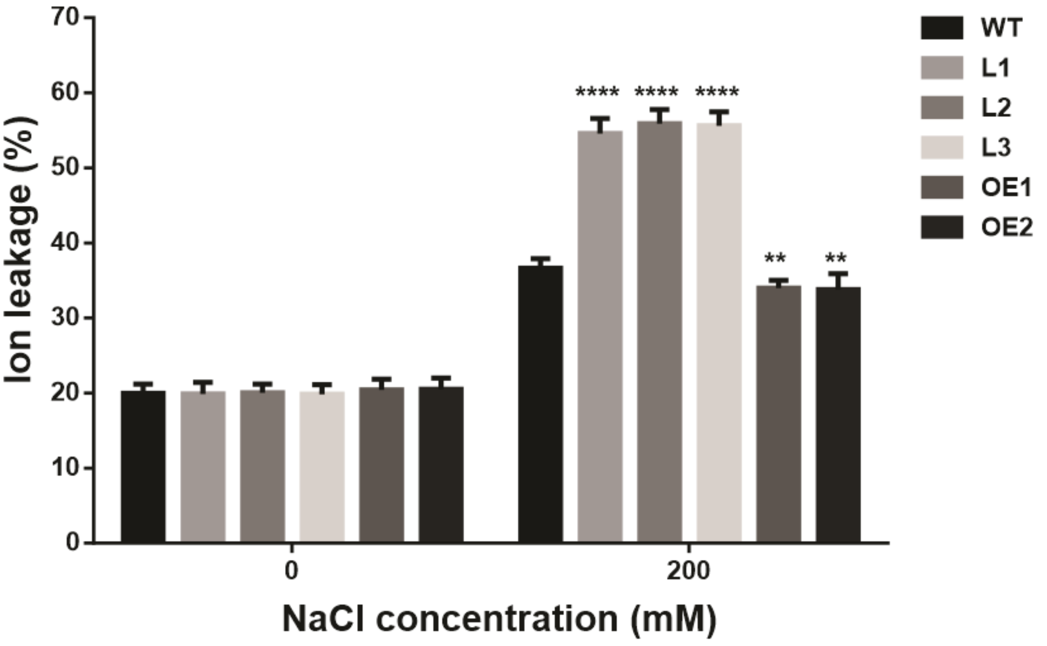


**Supplementary FIGURE S3.** Altered *VLG* expression affects membrane integrity under salt stress during long-day conditions. Ion leakage in WT, *VLG*-knock-down and *VLG*-overexpressing plants with or without salt treatment**.** Ion leakage was measured in leaves of four-week-old WT, *VLG*-knock-down (L1, L2, L3) and *VLG*-overexpressing (OE1, OE2) plants grown under long-day condition with or without 200 mM NaCl treatment for 3 weeks. ANOVA and Dunnett's multiple comparisons test $p<0.01$ (**), $p<0.0001$ (****). Three independent experiments were done with similar results.

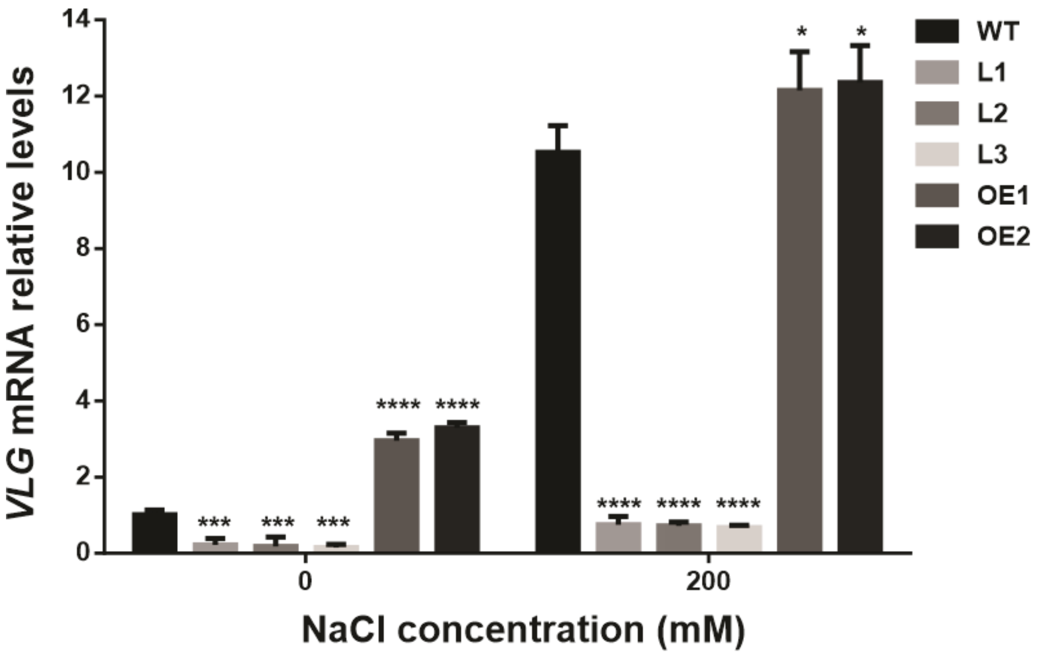


**Supplementary FIGURE S4.** Salt stress induces *VLG* expression in WT and *VLG*-overexpressing plants. Four-week-old WT, *VLG*-knock-down (L1, L2, L3), and *VLG*-overexpressing (OE1, OE2) plants were treated with or without 200 mM NaCl treatment for 3 days. The expression levels of *VLG* were measured by qRT-PCR. ANOVA and Dunnett's multiple comparisons test, $p<0.05$ (*), $p<0.001$ (***), $p<0.0001$ (****). Three independent experiments were done with similar results.

**Supplementary TABLE S1. Primers used for q-PCR**

| Primer | Sequence | Reference |
|---|---|---|
| *VLG* | F: 5′- GAAATGTGACTACGGTGCTCA - 3′ | (Amigo et al. 2025) |
| | R. 5′- CCACTTCCTCCTTCTCCTCCT - 3′ | |
| *ACT2* | F: 5′- GCCATCCAAGCTGTTCTCTC -3′ | (Soto et al. 2015) |
| | R: 5′- GAAACCCTCGTAGATTGGCA -3′ | |
| *SOS1* | F: 5′- TTGGCTATGGTTTGGATTGGA -3′ | (Song et al. 2023) |
| | R: 5′- ATGCGAAGAAGGCGTAGAACA -3′ | |
| *SOS3* | F: 5′- CATCCGTCACGCCATTCAC -3′ | (Song et al. 2023) |
| | R: 5′- TCACTCCATTTCGCTTCACATC -3′ | |
| *NHX1* | F: 5′- TTGAGCCTTCAGGGAACCA -3′ | (Song et al. 2023) |
| | R: 5′- AAAGCCACGACCTCCAAAGA -3′ | |
| *NHX3* | F: 5′- TGCCATAGGAACGATTTTCTCA -3′ | (Song et al. 2023) |
| | R: 5′- TCATTCACCACTCCTTCTCCAA -3′ | |
| *NHX5* | F: 5′- GCTGAAGGAGTTTCACAAAACCA -3′ | (Song et al. 2023) |
| | R: 5′- TCCCCATCTCCATCTCCATC -3′ | |
| *HKT1* | F: 5′- AAACGGCGAGAGATGTTCTTAGT -3′ | (Song et al. 2023) |
| | R: 5′- GGAGCCAGATGAGACCAGAGTT -3′ | |
| *RAP2.6* | F: 5′- AGTAAGGCAACGACCATGGG -3′ | (Song et al. 2023) |
| | R: 5′- GTCTCGAATGTCCCAAGCCA -3′ | |
| *RAP2.6L* | F: 5′- CAAGGCCCTACTACCACCACAA -3′ | (Krishnaswamy et al. 2011) |
| | R: 5′- GGTCGAGGAGGAGGTGAGTTC -3′ | |
| *LTL1* | F: 5′-GAACTGGTGCAATGGGATGT -3′ | This work |
| | R: 5′- TTCGATGTCACAAACCCAAA -3′ | |

**Supplementary TABLE 2. List of transcription factors present in the promoter region.**

| Gene Locus | TAIR ID | GO term | TAIR internal GO id | Reference |
|---|---|---|---|---|
| AT2G30250 | 2060805 | cellular response to heat | 29752 | PMID:21336597 |
| AT2G38340 | 2057217 | cellular response to heat | 29752 | PMID:21069430 |
| AT2G38470 | 2057212 | cellular response to heat | 29752 | PMID:21336597 |
| AT5G07100 | 2169354 | cellular response to heat | 29752 | PMID:21336597 |
| AT5G46350 | 2170403 | cellular response to hydrogen peroxide | 31391 | PMID:20367464 |
| AT1G01720 | 2198225 | cellular response to hypoxia | 33992 | PMID:31519798 |
| AT1G13260 | 2205319 | cellular response to hypoxia | 33992 | PMID:31519798 |
| AT1G25560 | 2031185 | cellular response to hypoxia | 33992 | PMID:31519798 |
| AT1G27730 | 2199246 | cellular response to hypoxia | 33992 | PMID:31519798 |
| AT2G46400 | 2039119 | cellular response to hypoxia | 33992 | PMID:31519798 |
| AT3G46080 | 2075276 | cellular response to hypoxia | 33992 | PMID:31519798 |
| AT3G46090 | 2075291 | cellular response to hypoxia | 33992 | PMID:31519798 |
| AT3G56400 | 2102539 | cellular response to hypoxia | 33992 | PMID:31519798 |
| AT4G01250 | 2125043 | cellular response to hypoxia | 33992 | PMID:31519798 |
| AT5G04340 | 2179964 | cellular response to hypoxia | 33992 | PMID:31519798 |
| AT5G05410 | 2153504 | cellular response to hypoxia | 33992 | PMID:31519798 |
| AT2G23320 | 2058568 | cellular response to oxygen-containing compound | 44644 | PMID:34562334 |
| AT5G19790 | 2183174 | cellular response to potassium ion | 38062 | PMID:22406475 |
| AT2G40340 | 2063073 | heat acclimation | 25138 | PMID:16807682 |
| AT3G11020 | 2085527 | heat acclimation | 25138 | PMID:16807682 |
| AT5G05410 | 2153504 | heat acclimation | 25138 | PMID:16807682 |
| AT5G43170 | 2167766 | hyperosmotic salinity response | 13163 | PMID:10806347 |
| AT5G62000 | 2174013 | positive regulation of potassium ion import across plasma membrane | 47956 | PMID:27895227 |
| AT5G46350 | 2170403 | positive regulation of response to salt stress | 41943 | PMID:23451802 |
| AT2G33860 | 2057609 | positive regulation of response to water deprivation | 45639 | PMID:29944444 |
| AT1G05805 | 505006103 | regulation of potassium ion import | 48481 | PMID:23779086 |
| AT1G51140 | 2026037 | regulation of potassium ion import | 48481 | PMID:23779086 |
| AT2G40750 | 2064806 | regulation of response to osmotic stress | 15450 | PMID:23815736 |
| AT3G56400 | 2102539 | regulation of response to osmotic stress | 15450 | PMID:23815736 |
| AT2G40750 | 2064806 | regulation of response to water deprivation | 35739 | PMID:28576847 |
| AT2G46400 | 2039119 | regulation of response to water deprivation | 35739 | PMID:28576847 |
| AT3G56400 | 2102539 | regulation of response to water deprivation | 35739 | PMID:28576847 |
| AT1G01060 | 2200970 | response to cold | 5433 | PMID:21471455 |
| AT1G27730 | 2199246 | response to cold | 5433 | PMID:15333755 |
| AT1G35460 | 2008693 | response to cold | 5433 | PMID:25246594 |
| AT1G49480 | 2010262 | response to cold | 5433 | PMID:22232549 |
| AT2G21060 | 2047092 | response to cold | 5433 | PMID:12529510 |
| AT2G30250 | 2060805 | response to cold | 5433 | PMID:18839316 |
| AT2G38470 | 2057212 | response to cold | 5433 | PMID:18839316 |
| AT2G46530 | 2039124 | response to cold | 5433 | PMID:34562334 |
| AT2G46590 | 2039959 | response to cold | 5433 | PMID:12084825 |
| AT2G46830 | 2044345 | response to cold | 5433 | PMID:21471455 |
| AT3G26744 | 2090847 | response to cold | 5433 | PMID:12672693 |

| | | | | |
|---|---|---|---|---|
| AT4G11070 | 2136093 | response to cold | 5433 | PMID:37622245 |
| AT4G25470 | 2131834 | response to cold | 5433 | PMID:16258011 |
| AT4G25470 | 2131834 | response to cold | 5433 | PMID:9952441 |
| AT4G25470 | 2131834 | response to cold | 5433 | PMID:9735350 |
| AT4G25480 | 2131849 | response to cold | 5433 | PMID:16258011 |
| AT4G25480 | 2131849 | response to cold | 5433 | PMID:9735350 |
| AT4G25480 | 2131849 | response to cold | 5433 | PMID:10096298 |
| AT4G25480 | 2131849 | response to cold | 5433 | PMID:12164808 |
| AT4G25490 | 2131854 | response to cold | 5433 | PMID:9023378 |
| AT4G25490 | 2131854 | response to cold | 5433 | PMID:9707537 |
| AT4G25490 | 2131854 | response to cold | 5433 | PMID:16258011 |
| AT4G25490 | 2131854 | response to cold | 5433 | PMID:9735350 |
| AT4G26440 | 2131503 | response to cold | 5433 | PMID:20643804 |
| AT4G38680 | 2121219 | response to cold | 5433 | PMID:12529510 |
| AT4G38680 | 2121219 | response to cold | 5433 | PMID:17169986 |
| AT4G38680 | 2121219 | response to cold | 5433 | PMID:17963727 |
| AT5G43170 | 2167766 | response to cold | 5433 | PMID:10806347 |
| AT3G04070 | 2095908 | response to flooding | 5789 | PMID:24363315 |
| AT1G12610 | 2195052 | response to freezing | 18008 | PMID:21421412 |
| AT3G26744 | 2090847 | response to freezing | 18008 | PMID:17416732 |
| AT1G12610 | 2195052 | response to heat | 5962 | PMID:21421412 |
| AT1G30650 | 2204549 | response to heat | 5962 | PMID:36155833 |
| AT1G35460 | 2008693 | response to heat | 5962 | PMID:25246594 |
| AT2G38470 | 2057212 | response to heat | 5962 | PMID:18839316 |
| AT2G38880 | 2064996 | response to heat | 5962 | PMID:25490919 |
| AT2G46830 | 2044345 | response to heat | 5962 | PMID:25246594 |
| AT3G05690 | 2078072 | response to heat | 5962 | PMID:25490919 |
| AT4G14540 | 2129885 | response to heat | 5962 | PMID:25490919 |
| AT4G14540 | 2129885 | response to heat | 5962 | PMID:31123093 |
| AT4G14540 | 2129885 | response to heat | 5962 | PMID:25490919 |
| AT5G05410 | 2153504 | response to heat | 5962 | PMID:17030801 |
| AT5G05410 | 2153504 | response to heat | 5962 | PMID:25490919 |
| AT5G67300 | 2158212 | response to heat | 5962 | PMID:31358650 |
| AT3G29035 | 2087037 | response to hydrogen peroxide | 13166 | PMID:21303842 |
| AT4G23810 | 2128514 | response to hydrogen peroxide | 13166 | PMID:22268143 |
| AT5G05410 | 2153504 | response to hydrogen peroxide | 13166 | PMID:17030801 |
| AT5G24110 | 2167428 | response to hydrogen peroxide | 13166 | PMID:22268143 |
| AT5G39610 | 2164895 | response to hydrogen peroxide | 13166 | PMID:20404534 |
| AT3G14230 | 2090975 | response to hypoxia | 10824 | PMID:20357136 |
| AT3G20770 | 2091906 | response to hypoxia | 10824 | PMID:25284079 |
| AT1G51140 | 2026037 | response to osmotic stress | 6618 | PMID:24261563 |
| AT1G51140 | 2026037 | response to osmotic stress | 6618 | PMID:24261563 |
| AT1G66550 | 2028962 | response to osmotic stress | 6618 | PMID:34562334 |
| AT1G69310 | 2007081 | response to osmotic stress | 6618 | PMID:22930734 |
| AT2G30250 | 2060805 | response to osmotic stress | 6618 | PMID:18839316 |
| AT2G38470 | 2057212 | response to osmotic stress | 6618 | PMID:18839316 |
| AT2G40750 | 2064806 | response to osmotic stress | 6618 | PMID:23815736 |
| AT3G56400 | 2102539 | response to osmotic stress | 6618 | PMID:23815736 |
| AT1G27730 | 2199246 | response to oxidative stress | 6625 | PMID:18156220 |

| | | | | |
|---|---|---|---|---|
| AT1G66550 | 2028962 | response to oxidative stress | 6625 | PMID:34562334 |
| AT2G23320 | 2058568 | response to oxidative stress | 6625 | PMID:34562334 |
| AT3G46090 | 2075291 | response to oxidative stress | 6625 | PMID:14722088 |
| AT5G18270 | 2172334 | response to oxidative stress | 6625 | PMID:34562334 |
| AT5G39610 | 2164895 | response to oxidative stress | 6625 | PMID:15295076 |
| AT2G40750 | 2064806 | response to reactive oxygen species | 10197 | PMID:22268143 |
| AT3G56400 | 2102539 | response to reactive oxygen species | 10197 | PMID:22268143 |
| AT4G17490 | 2129106 | response to reactive oxygen species | 10197 | PMID:23300166 |
| AT5G19790 | 2183174 | response to reactive oxygen species | 10197 | PMID:22406475 |
| AT1G17950 | 2030903 | response to salt | 45327 | PMID:21399993 |
| AT1G17950 | 2030903 | response to salt | 45327 | PMID:21399993 |
| AT1G51140 | 2026037 | response to salt | 45327 | PMID:24261563 |
| AT1G51140 | 2026037 | response to salt | 45327 | PMID:24261563 |
| AT5G39610 | 2164895 | response to salt | 45327 | PMID:20404534 |
| AT1G12610 | 2195052 | response to salt stress | 7182 | PMID:18643985 |
| AT1G13960 | 2014799 | response to salt stress | 7182 | PMID:34351541 |
| AT1G27730 | 2199246 | response to salt stress | 7182 | PMID:8662738 |
| AT1G27730 | 2199246 | response to salt stress | 7182 | PMID:8662738 |
| AT1G69310 | 2007081 | response to salt stress | 7182 | PMID:22930734 |
| AT1G80590 | 2198963 | response to salt stress | 7182 | PMID:36834483 |
| AT2G03340 | 2063835 | response to salt stress | 7182 | PMID:34351541 |
| AT2G23320 | 2058568 | response to salt stress | 7182 | PMID:34562334 |
| AT2G30250 | 2060805 | response to salt stress | 7182 | PMID:18839316 |
| AT2G38340 | 2057217 | response to salt stress | 7182 | PMID:21069430 |
| AT2G38470 | 2057212 | response to salt stress | 7182 | PMID:18839316 |
| AT2G41835 | 504955994 | response to salt stress | 7182 | PMID:34562334 |
| AT2G42400 | 2053786 | response to salt stress | 7182 | PMID:30477148 |
| AT3G01470 | 2084228 | response to salt stress | 7182 | PMID:16055682 |
| AT3G61830 | 2076765 | response to salt stress | 7182 | PMID:34562334 |
| AT5G18270 | 2172334 | response to salt stress | 7182 | PMID:34562334 |
| AT5G39610 | 2164895 | response to salt stress | 7182 | PMID:16359384 |
| AT5G39610 | 2164895 | response to salt stress | 7182 | PMID:20113437 |
| AT5G39610 | 2164895 | response to salt stress | 7182 | PMID:19608714 |
| AT5G67300 | 2158212 | response to salt stress | 7182 | PMID:18162593 |
| AT1G12610 | 2195052 | response to water deprivation | 5647 | PMID:21421412 |
| AT1G17950 | 2030903 | response to water deprivation | 5647 | PMID:21399993 |
| AT1G27730 | 2199246 | response to water deprivation | 5647 | PMID:15333755 |
| AT1G27730 | 2199246 | response to water deprivation | 5647 | PMID:15333755 |
| AT1G33240 | 2196663 | response to water deprivation | 5647 | PMID:21169508 |
| AT1G51140 | 2026037 | response to water deprivation | 5647 | PMID:24261563 |
| AT1G51140 | 2026037 | response to water deprivation | 5647 | PMID:24261563 |
| AT1G52890 | 2011531 | response to water deprivation | 5647 | PMID:15319476 |
| AT1G54160 | 2014375 | response to water deprivation | 5647 | PMID:18682547 |
| AT1G69310 | 2007081 | response to water deprivation | 5647 | PMID:22930734 |
| AT2G23320 | 2058568 | response to water deprivation | 5647 | PMID:34562334 |
| AT2G38340 | 2057217 | response to water deprivation | 5647 | PMID:21069430 |
| AT2G38470 | 2057212 | response to water deprivation | 5647 | PMID:18839316 |
| AT2G38880 | 2064996 | response to water deprivation | 5647 | PMID:25490919 |
| AT2G38880 | 2064996 | response to water deprivation | 5647 | PMID:17923671 |

| | | | | |
|---|---|---|---|---|
| AT3G11020 | 2085527 | response to water deprivation | 5647 | PMID:10809011 |
| AT3G15500 | 2090176 | response to water deprivation | 5647 | PMID:15319476 |
| AT3G58710 | 2099044 | response to water deprivation | 5647 | PMID:34562334 |
| AT3G61830 | 2076765 | response to water deprivation | 5647 | PMID:34562334 |
| AT4G14540 | 2129885 | response to water deprivation | 5647 | PMID:25490919 |
| AT4G14540 | 2129885 | response to water deprivation | 5647 | PMID:31123093 |
| AT4G25480 | 2131849 | response to water deprivation | 5647 | PMID:11158529 |
| AT4G25480 | 2131849 | response to water deprivation | 5647 | PMID:16617101 |
| AT4G25490 | 2131854 | response to water deprivation | 5647 | PMID:9023378 |
| AT4G27410 | 2124014 | response to water deprivation | 5647 | PMID:15319476 |
| AT4G27410 | 2124014 | response to water deprivation | 5647 | PMID:15341629 |
| AT4G38680 | 2121219 | response to water deprivation | 5647 | PMID:17169986 |
| AT5G05410 | 2153504 | response to water deprivation | 5647 | PMID:10809011 |
| AT5G05410 | 2153504 | response to water deprivation | 5647 | PMID:16617101 |
| AT5G07680 | 2160324 | response to water deprivation | 5647 | PMID:34562334 |
| AT5G18270 | 2172334 | response to water deprivation | 5647 | PMID:34562334 |
| AT5G47640 | 2168983 | response to water deprivation | 5647 | PMID:25490919 |
| AT5G47670 | 2169028 | response to water deprivation | 5647 | PMID:25490919 |
| AT5G61590 | 2151576 | response to water deprivation | 5647 | PMID:18552355 |
| AT5G67300 | 2158212 | response to water deprivation | 5647 | PMID:18552355 |
